# Chromosome-scale genome assembly of the rough periwinkle *Littorina saxatilis*

**DOI:** 10.1101/2024.02.01.578480

**Authors:** Aurélien De Jode, Rui Faria, Giulio Formenti, Ying Sims, Timothy P. Smith, Alan Tracey, Jonathan M. D. Wood, Zuzanna B. Zagrodzka, Kerstin Johannesson, Roger K. Butlin, Erica H Leder

## Abstract

The intertidal gastropod *Littorina saxatilis* is a model system to study speciation and local adaptation. The repeated occurrence of distinct ecotypes showing different levels of genetic divergence makes *L. saxatilis* particularly suited to study different stages of the speciation continuum in the same lineage. A major finding is the presence of several large chromosomal inversions associated with the divergence of ecotypes and, specifically, the species offers a system to study the role of inversions in this divergence. The genome of *L. saxatilis* is 1.35Gb and composed of 17 chromosomes. The first reference genome of the species was assembled using Illumina data, was highly fragmented (N50 of 44kb) and was quite incomplete, with a BUSCO completeness of 80.1% on the Metazoan dataset. A linkage map of one full-sibling family enabled the placement of 587 Mbp of the genome into 17 linkage groups corresponding to the haploid number of chromosomes, but the fragmented nature of this reference genome limited the understanding of the interplay between divergent selection and gene flow during ecotype formation. Here we present a newly generated reference genome that is highly contiguous, with a N50 of 67 Mb and 90.4% of the total assembly length placed in 17 super-scaffolds. It is also highly complete with a BUSCO completeness of 94.1 % of the Metazoa dataset. This new reference will allow for investigations into the genomic regions implicated in ecotype formation as well as better characterization of the inversions and their role in speciation.

**Significance:** The rough periwinkle, *L. saxatilis* has become a model to study adaptation, evolutionary innovation and speciation, including the role of chromosomal inversions in these processes. Chromosomal inversions have also been identified in other species of *Littorina* providing a valuable opportunity for investigating their origin and evolution. Here, we present a new chromosome-scale reference genome of *L. saxatilis* generated from long reads and HiC chromatin proximity mapping replacing an earlier highly fragmented genome. This will enable detailed investigations of how inversions contribute to adaptation and reproductive isolation in *L. saxatilis* and related taxa, including studies of the effects of individual loci within and outside inversions.

## Introduction

Speciation and local adaptation are key evolutionary processes that have been successfully studied in the marine snail, *Littorina saxatilis* (Figure 1A) and related taxa (Blakeslee et al. 2021; Johannesson 2016; Johannesson et al., 2010; 2017; Rolán-Alvarez et al., 2015). Multiple ecotypes have been described in this species and its close relatives (Reid 1996; Johannesson et al 2023). In particular, two ecotypes are repeatedly found in close proximity on the North Atlantic shores from Portugal to Norway and have been studied in detail. The ‘crab’ ecotype is typically found in areas with intense crab predation while the ‘wave’ ecotype is present in shore areas battered by waves. Their distributions often partially overlap in contact zones. The two ecotypes differ in several traits such as shell size, shape, thickness, ornamentation and behaviour. Across the species distribution, gene flow is occurring at different rates between the two ecotypes. In Sweden, allelic frequencies present a clinal pattern from crab to wave habitat (Westram et al. 2021), while in Spain two distinct genetic clusters are found in upper and lower shore areas (Raffini et al. 2023). This particular configuration enables the characterization of genomic, phenotypic, and organismal differences between pairs of populations from the same species at various stages of divergence, making *L. saxatilis* a most appropriate system to study the speciation process (Johannesson et al. 2023). Ecotypes also form in other species, providing opportunities for comparative studies of divergence mechanisms for population pairs along the speciation continuum (Johannesson et al. 2023).

**Figure 1.**
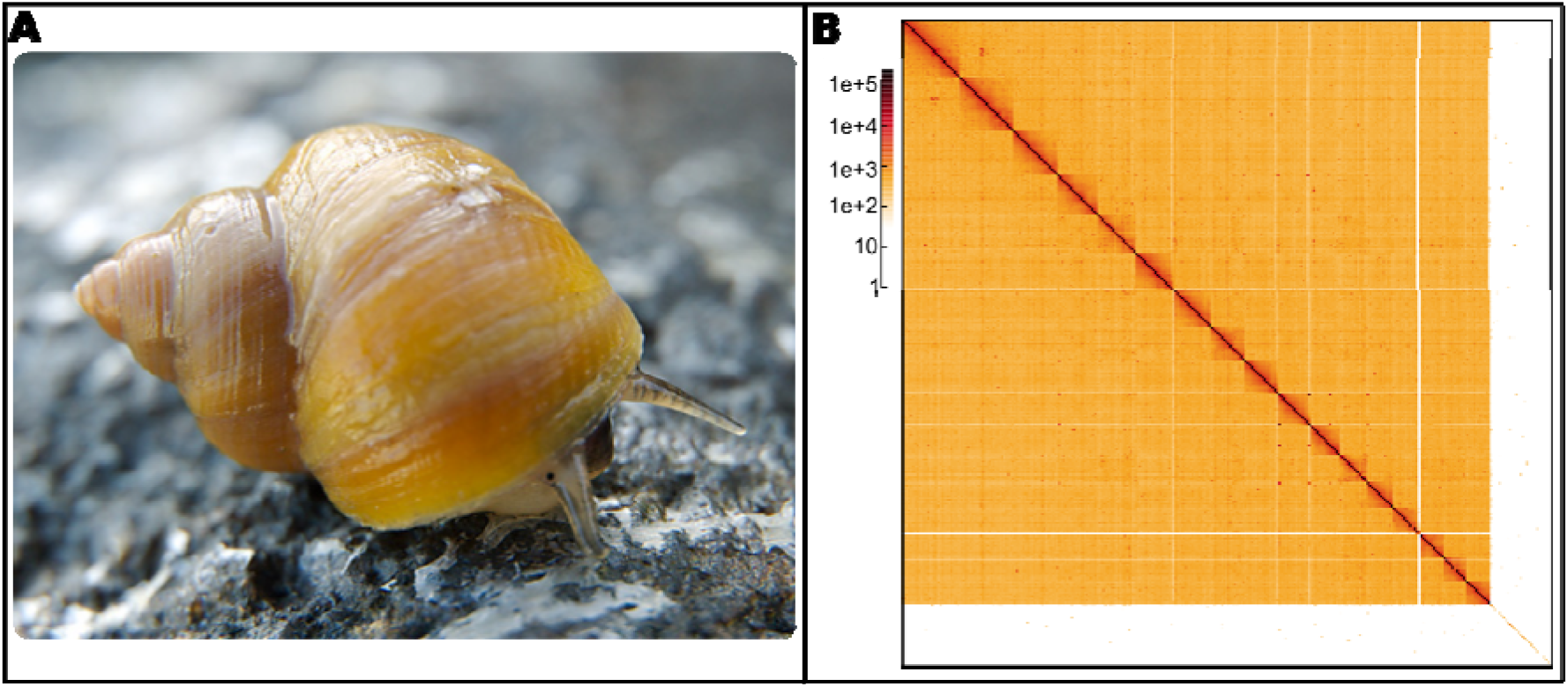
*Littorina saxatilis* crab ecotype picture and Hi-C map. A. Picture of *Littorina saxatilis* crab ecotype on Swedish shore. B. Hi-C contact map of the new *Littorina saxatilis* assembly after manual curation, visualised in HiGlass. Chromosomes are arranged in size order from left to right and top to bottom.

Moreover, several genomic inversions showing allelic frequency differences between ecotypes (Faria et al. 2019a; Westram et al. 2021), sex association (Hearn et al. 2022) and association with adaptive traits (Koch et al. 2021; 2022) have been identified in *L. saxatilis* and other species of the genus (Reeve et al. 2023; Le Moan et al. 2023). Inversions can promote the process of local adaptation and speciation and are associated with ecotype divergence in several other natural systems, *e*.*g*. (Faria et al. 2019b; Merot 2020).

Furthermore, *Littorina saxatilis* and its near relatives have direct development, *i*.*e*., no planktonic stage (Reid 1996), resulting in low dispersal; however, *L. saxatilis* is ovoviviparous while the other species lay eggs on the substrate. The shift in reproductive mode compared to other members of the genus provides an outstanding opportunity to study the evolution of this key innovation (Stankowski et al. 2023).

Therefore, *L. saxatilis* and the whole *Littorina* genus constitute a particularly interesting system to study the importance of chromosomal inversions in local adaptation and reproductive isolation. However, for this, a contiguous reference genome is needed.

The *L. saxatilis* genome is composed of 17 chromosomes for a total haploid size of around 1.35 Gbp (Birstein and Mikhailova, 1990; Janson 1983; Rolán[Alvarez et al. 1996). The first reference genome of *L. saxatilis* (hereafter “*L. saxatilis* genome v.1”) was 1.6 Gbp in length and composed of 388,619 contigs placed in 116,262 scaffolds (Westram et al. 2018). This genome was highly fragmented with a N50 of ∼44 kbp, with only 10% of its contigs (representing 50% of its total length) placed into 17 linkage groups by linkage mapping, and a BUSCO completeness of 80.1% against the Metazoa reference (Westram et al. 2018). This has constituted an obstacle to the precise characterization of the genomic architecture of, for example, the divergence between ecotypes. Such characterisation is necessary to properly address questions regarding speciation, local adaptation, and to achieve a detailed understanding of the involvement of inversions in these processes.

Here we present the first chromosome scale, annotated genome of *L. saxatilis* (hereafter “*L. saxatilis* genome v.2”). This new reference constitutes a major improvement compared to the existing one and will enable us to further enhance our understanding of evolution in this system.

## Results and Discussion

A total of 24,891,710 PacBio CLR reads were generated representing 174,323,538,425 base pairs. Assembling with CANU (Koren et al. 2017) produced 23,755 contigs for a total length of 2,278,220,962 bases. After two rounds of haplotig removal with purge_dups and long read polishing, the assembly contained 9,667 contigs for a total length of 1,261,606,186 bases. While BUSCO (Manni et al. 2021) completeness remained at 95.2% for the Metazoa dataset after the two rounds of haplotig removal with purge_dups, the ‘complete and duplicated’ score went from 72% to 2.9%. Merqury (Rhie et al. 2020) copy number spectra (spectra_cn) plots also showed a clear decrease in the amount of haplotypic duplication retained in the assembly (Fig. S1). Some level of haplotypic duplication could still be seen on the spectra graphs, but neither an additional round of purge_dups nor parameter tuning could remove the remaining haplotypic duplication. The high levels of haplotypic duplication obtained in the raw assembly are most likely due to the high heterozygosity of our genome (∼1.5% based on k-mer analysis using GenomeScope (Vurture et al. 2017)) combined with potential polymorphic inversions detected in the *L. saxatilis* genome being heterokaryotypic in the individual we sequenced. The final k-mer spectra (Fig. S1) showed a high number of k-mers only found in the assembly and a high frequency of k-mers found only in the reads set (peak of the black line around 30X). This is most likely due to the fact that the Illumina reads used for the kmer analysis came from a different individual than the individual used to assemble the genome. Using Blobtoolkit (Laetsch and Blaxter 2017; Challis et al. 2020), a total of 55 contigs identified as bacteria or Ciliophora (known commensals of *L. saxatilis* (Fenchel 1965)) were removed from the assembly prior to scaffolding.

A total of 280 million reads was obtained for Hi-C sequencing. The scaffolding produced a total of 4,322 scaffolds, and the scaffolded assembly had a N50 of around 82 Mb. Several errors made by the automatic scaffolding procedure implemented in YaHS (Zhou et al. 2023) were corrected by manual curation (Figure 1B, Fig. S2). The final assembly was composed of 4,094 scaffolds including 17 super-scaffolds (composed of contigs joined by gap with a minimum arbitrary size of 200 bp) that contained 90.4 % of the total base pairs in the assembly (Figure 2A, Fig. S3). The hexamer telomere motif of TTAGGG was found on 16 scaffolds but given the noisy nature of CLR long reads, the assembly of these regions is likely inaccurate and under-representing the length of these regions. The generated assembly was highly contiguous with an N50 of 67 Mb and a completeness BUSCO score of 98.4 %, 94.1 % and 79.2 % for the Eukaryota, Metazoa and Mollusca databases, respectively (Figure 2B, Table S1). The 17 super-scaffolds corresponded to the number of expected chromosomes for *L. saxatilis* (Birstein and Mikhailova, 1990, Janson 1983, Rolán□Alvarez, Buño, & Gosalvez, 1996, Faria et al. 2019a) and were matched to the linkage groups (LG) from the *L. saxatilis* genome v.1 (Table S2). Using RepeatMasker (v 4.1.5) in quick search mode, simple repeats were identified comprising 7.81 % of the genome, while low complexity regions comprised 0.89% of the genome. Additionally, 199 small RNAs were identified.

**Figure 2.**
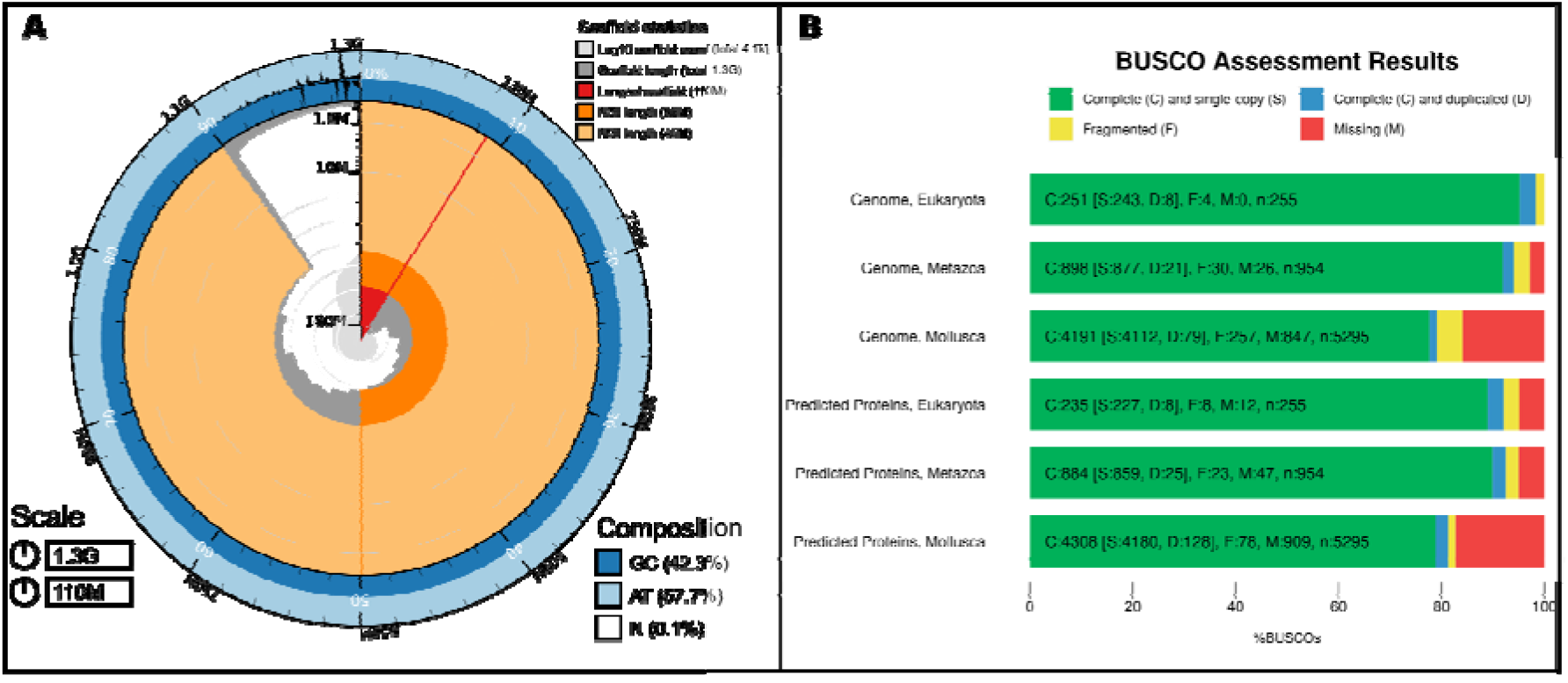
Genome assembly of *Littorina saxatilis*: metrics. A. BlobToolKit Snailplot shows N50 metrics. The main plot is divided into 1,000 size-ordered bins around the circumference with each bin representing 0.1% of the 1,256,336,603 bp assembly. The distribution of scaffold lengths is shown in dark grey with the plot radius scaled to the longest scaffold present in the assembly (111,777,080 bp, shown in red). Orange and pale-orange arcs show the N50 and N90 scaffold lengths (67,515,031 and 44,900,717 bp), respectively. The pale grey spiral shows the cumulative scaffold count on a log scale with white scale lines showing successive orders of magnitude. The blue and pale-blue area around the outside of the plot shows the distribution of GC, AT and N percentages in the same bins as the inner plot. B. Busco scores for the genome and gene sets on the Metazoa and Mollusca datasets.

There were 25,317 protein coding genes predicted by the Braker3 pipeline (Gabriel et al. 2023). The majority of these genes had only one protein predicted, but 3,486 had two or more protein variants. The number of predicted proteins is slightly higher, but still quite similar to other gastropods: 21,438 *Gigantopelta aegis* (Lan et al. 2021), 21,533 *Pomacea canaliculata* (Liu et al. 2018), 23,800 *Lottia gigantea* (Simakov et al. 2013). BUSCO scores of completeness for the predicted proteins were 93.0% and 82.2 % for the Metazoa and Mollusca databases, respectively (Figure 2B, Table S1).

A mitochondrial genome of *L. saxatilis* was previously assembled using the Illumina data (Marques et al. 2017) and is available on Genbank (NC_030595.1).

### Conclusion

We combined PacBio CLR reads and Hi-C chromatin proximity mapping data to generate the first chromosome-level genome assembly for *Littorina saxatilis*. The *L. saxatilis* genome v.2 achieved a scaffold N50 of 67Mb with more than 90% of the total assembly length (1.2Gb) placed in 17 super-scaffolds. Moreover, this genome is highly contiguous with a 98% completeness BUSCO score on the Eukaryota dataset and contained 25,317 protein coding genes. The *L. saxatilis* genome v.2 constitutes a major improvement compared to the *L. saxatilis* genome v.1 and will underpin significant advances in the study of speciation and evolution, specifically the evolution of chromosomal inversions and their role in diversification within the *Littorina* genus.

## Material and Methods

High Molecular Weight DNA was extracted from fresh head and foot tissue of one female ‘crab’ ecotype of *L. saxatilis* collected in Sweden (N58°52’28”, E11°6’59”) following the protocol developed by Grohme et al. (2018) with minor adjustments. The protocol consisted of three parts: mucus removal using N-acetyl-L-cysteine, gDNA isolation using a phenol–chloroform-isoamyl alcohol solution and post purification of gDNA using CTAB (see full protocol in Supp.). The library was prepared with PacBio’s SMRTbell Express Template Prep Kit 2.0, and sequenced using the Sequel Binding Kit 3.0, Sequel Sequencing Plate 3.0 and Sequel DNA Internal Control 3.0. A total of 16 PacBio ZMW cells were sequenced using a Sequel instrument. Raw PacBio BAM files were converted to fastq files using bam2fastq v1.0.0 (https://github.com/jts/bam2fastq) from the SMRT suite. CLR PacBio raw reads were assembled using CANU v2.0 (Koren et al. 2017) with default parameters. Haplotypic duplications were removed from the raw assembly using the purge_dups v1.2.5 (Guan et al. 2020) pipeline two times in a row with default parameters. We used the short Illumina reads that were assembled to generate *L. saxatilis* genome v.1 (Westram et al. 2018) in the Merqury, Merfin and Nextpolish analysis described hereafter. Levels of haplotypic duplication at the different steps were assessed using spectra-cn plot from Merqury (Rhie et al. 2020) based on Illumina reads. The obtained genome was polished first using the raw CLR long reads with the arrow_grid wrapper (Chin et al. 2013; Koren et al. 2017). The vcf file obtained was corrected using Merfin v1.1-development (Formenti at al. 2022) based on Illumina reads. This polishing procedure was run 2 times. Finally, raw Illumina reads were trimmed and filtered using fastp (Chen et al. 2018) with the following parameters --cut_front -- cut_front_window_size 1 --cut_front_mean_quality 30 --cut_right --cut_right_mean_quality 3. The obtained quality filtered reads were used in a final round of polishing implemented in NextPolish v1.3.1 (Hu et al. 2020). Prior to scaffolding, contigs were examined for contaminants using the Blobtoolkit v2.6.4 pipeline (Laetsch and Blaxter 2017; Challis et al. 2020) and 55 of the contigs were removed from the assembly.

A different ‘crab’ ecotype female snail from the same location Sweden was used to build a Dovetail Hi-C library using the DpnII enzyme. Paired end reads (2×150 bp) were generated at SciLife on one lane of a NovaSeq 6000 SP instrument. Raw Hi-C reads were mapped to the contigs following the Arima mapping pipeline (https://github.com/ArimaGenomics/mapping_pipeline/tree/master), and contigs were placed into scaffolds using YaHS v1.1a-r3 (Zhou et al. 2023). The obtained scaffolded genome was further manually curated (Howe et al. 2021) using pretext (https://github.com/wtsi-hpag/PretextView), and HiGlass (Kerpedjiev et al. 2018).

Finally, assembly contiguity and completeness were assessed using gfastats v1.3.6 v5.0.2 (Formenti et al. 2022), BUSCO v5.4.7 (Manni et al. 2021) and the Blobtoolkit v2.6.4 pipeline (Laetsch and Blaxter 2017; Challis et al. 2020).

To establish correspondence between the super-scaffolds from *L. saxatilis* genome v.1 and the linkage groups of *L. saxatilis* genome v.2 the following procedure was followed. For each SNP placed in the linkage map from the *L. saxatilis* genome v.1 a 200 bp sequence centered on the SNP was extracted and mapped to the *L. saxatilis* genome v.2 using minimap2 (Li 2018). Alignments were filtered by removing duplicates and partial mappings that did not include the SNP. Then for each 200 bp fragment, the best alignment was selected. RepeatMasker (v 4.1.5) was used to identify simple repeats using the quick search option (http://www.repeatmasker.org).

RNAseq data from Genbank BioProject PRJNA550990 along with RNAseq from mantle tissue from three ‘crab’ ecotype females from Sweden were mapped to the genome assembly using HiSat2 v2.2.1 (Kim et al. 2019). Reads were sorted and filtered to include only mapped reads using SAMtools v 1.16.1 (Li et al. 2009). BRAKER3 (Gabriel et al. 2023) and the algorithms therein (Brůna et al. 2021, Gotoh 2008, Hoff et al. 2016, Hoff et al. 2019, Iwata & Gotoh, 2012, Stanke et al. 2006, and Stanke et al. 2008) were used to predict protein coding genes in the assembly using both the RNAseq data from *Littorina saxatilis* and mollusc proteins from OrthoDBv11 (Kuznetsov et al. 2023). In addition to the algorithms employed by BRAKER3, the pipeline also uses various tools to gather evidence for protein sequences from the RNAseq data and the protein data using the following tools: DIAMOND (Buchfink et al. 2015), Stringtie2 (Kovaka et al. 2019), GFF utilities (Pertea & Pertea, 2020), BamTools (Barnett et al. 2011), and BEDTools (Quinlan, 2014). The protein evidence obtained was used for training GeneMark-EPT (Brůna et al. 2023) and later AUGUSTUS (Stanke et al. 2006), and then the two sets of predictions were combined using TSEBRA (Gabriel et al. 2023). The Braker3 Docker container version v.1.0.4 was used for this analysis. BLASTP was then used to identify the predicted proteins using a custom database of proteins from *Crassotrea virginica, Lottia gigantea, Gigantopelta aegis*, and *Pomacea canaliculata*.

## Acknowledgments

*The computations and data handling were enabled by resources in project NAISS 2023/6-270 provided by the National Academic Infrastructure for Supercomputing in Sweden (NAISS) at UPPMAX, funded by the Swedish Research Council through grant agreement no. 2022-06725*. Sequencing was performed at the Genomics Laboratory Facility, School of Biosciences, University of Sheffield. We thank Rachel Tucker and Helen Hipperson for their valuable input. The computations for the annotation were performed on resources provided by Sigma2 - the National Infrastructure for High Performance Computing and Data Storage in Norway. We are very grateful to Sergey Koren for his help in getting CANU running on the slurm cluster. We also thank Alan Le Moan for proving the correspondence table between *L. saxatilis* genome v1 and *L. saxatilis* genome v2. This work was funded by the European Research Council (ERC-2015-AdG-693030-BARRIERS, RKB), and the Swedish Research Council VR (2018-03695, RKB; 2021-04191, KJ; 2020-05385, EHL). RF was funded by the European Union’s Horizon 2020 research and innovation programme under the Marie Sklodowska-Curie (grant agreement No. 706376) and by the Fundação para a Ciência e a Tecnologia (2020.00275.CEECIND and PTDC/BIA-EVL/1614/2021). The use of trade names or commercial products in this manuscript is solely to provide specific information. It does not imply recommendation or endorsement by the U.S. Department of Agriculture. The USDA is an equal opportunity provider and employer.

## Data Availability

The PacBio CLR sequencing reads, the sequences from the Hi-C library and the Illumina short reads are deposited in NCBI under accession number PRJNA850123. The transcriptome short-read sequences are deposited under accession number PRJNA550990. The final genome assembly and genome annotation have been deposited in Genbank accession number -------.

