## Supplementary info for "Chromosome-scale genome assembly of the rough periwinkle *Littorina saxatilis*"

Contents

**DNA extraction protocol** .......................................................…...............….......................Page 2-4

**k-mer spectra plots** ...............................................................................................................Page 5

**Correspondence between *L. saxatilis* genome v1 and *L. saxatilis* genome v2.......…..**...Page 6

**Detailed DNA extraction protocol**

From: DNA extraction protocol: Small- and Large-Scale High Molecular Weight Genomic DNA Extraction from Planarians by Grohme et al. (2018) - **DOI:** 10.1007/978-1-4939-7802-1_7

Modified by Zuzanna Zagrodzka (University of Sheffield) for *Littorina saxatilis* tissue

1. Mucus Removal from Planarian Surface

1. For 10 ml of NAC mucus stripping solution, weigh 50 mg of NAC into a 15 ml tube and fill the tube to 8 ml with

ddH2O. This is typically sufficient for 10× 10 mm animals.

2. Add 10 μl of Phenol Red, the solution should turn bright yellow.

3. Add 200 μl 1 M HEPES-NaOH (pH 7.25– 7.4) and mix well.

4. With intermittent mixing, slowly add ~300–350 μl 1 M NaOH to the solution while monitoring the colour. A faint

red colour indicates a pH >7. A bright magenta colouring indicates alkaline pH around 8 and higher. In this case,

repeat the whole procedure.

Note: NAC is acidic in aqueous solution and the highest mucolytic activity is observed between pH 7 and 9.

4. Fill the tube up to 10 ml with distilled H2O.

5. Cut tissue (head and foot) into smaller pieces using sterile scalpel and put for 30 min into the solution at room

temperature with strong agitation (e.g., rotator).

6. Wash the tissue briefly with distilled water and process them further. This protocol is a single tube lysis and

organic.

2. HMW DNA Isolation Protocol for Cloning and Next Generation Sequencing

Carry out all procedures at room temperature (RT) unless otherwise specified. Perform steps 2–8 in the fume hood,

except for the digestion step (2) but be sure that tubes are sealed tightly. Use only wide-bore tips and mix the

samples only by careful inversion. Typically, the extracted DNA runs at a size of 50–400 kbp on a pulse field

electrophoresis gel.

1. In the fume hood add 500 μl of cold GTC buffer into a 1.5 ml tube. If you didn’t add mercaptoethanol earlier, add

now 3.5 ul of mercaptoethanol and mix well.

2. Place a tissue of ~3 mm into the tube. Take a small strip of Parafilm and stretch around the edges to seal the tube.

Incubate at 4oC (in a walk-in fridge) on a tube rotator for 2 hours. The tissue should dissociate entirely and no tissue

pieces should be visible anymore.

Note: Be careful not to exceed maximal centrifugation speed for the respective vessels.

3. Add 1 vol of phenol–chloroform-isoamyl alcohol to the lysate. Mix slowly by inversion for 10 minutes.

4. Centrifuge for 5 min at 10,000 × g at 4 °C to separate the phases.

5. Using a pipette with wide-bore tips, carefully transfer the upper aqueous phase to a new tube without disturbing

the interphase.

Note: For high molecular weight DNA the usage of wide bore or cut off pipette tips is advised.

6. Add 1 vol of chloroform to the aqueous phase and mix by inversion for 10 minutes to remove residual phenol.

7. Centrifuge for 5 min at 10,000 × g at 4 °C to separate the phases. Transfer the upper aqueous phase to a new tube.

8. Add 1 vol of cold 5 M NaCl and mix well by inverting the tube several times. The solution should turn cloudy for

concentrated samples. Place tube on ice for 15 min to salt out the contaminants.

9. Spin tube for 15 min at 12,000–16,000 × g in a centrifuge at 4 °C (from this step on you don’t have to work under

the fume hood).

10. Carefully decant or pipet off the supernatant containing the nucleic acids without disturbing the pellet to a fresh

2 ml tube.

11. Add 1 vol of isopropanol to the supernatant. Invert several times to mix. If you can’t see loads of DNA proceed to

the step no 13. If you can see a fluff of precipitated DNA retrieve it with a closed Pasteur pipette (glass rod) by

spinning the rod in the fluff. Rinse the DNA on the rod in 70% ethanol two times (prepare two separate tubes with

EtOH and spin the rod with DNA gently in each of the tubes. Dry the DNA fluff in air on the rod for a few minutes

(until the ethanol evaporates).

12. Dissolve in 50 μl of appropriate buffer, preferably AE buffer (low TE). Let the rod to stay in the tube and let the

DNA dissolve overnight in the room temperature. Proceed to: Postpurification of DNA Using CTAB.

13. Incubate 15–30 min on ice.

14. Spin at 2000–4000 × g for 30–45 min at 25 °C for HMW DNA.

15. Carefully decant or pipet off the supernatant without disturbing the pellet.

14. Wash the pellet by adding 1 ml of 70% ethanol to the tube and invert for 15 minutes. Make sure to dislodge the

pellet to remove the salt from it.

15. Spin the tube for an additional 5 min.

Note: Caution: the pellet might dislodge easily after the ethanol wash.

16. Let the pellet resuspend in 50 μl of appropriate buffer, preferably AE buffer (low TE) overnight in the room

temperature.

Note: At this point you have DNA and coisolated RNA in your sample.

3. Postpurification of DNA Using CTAB

Carry out all procedures at room temperature as CTAB precipitates at temperatures lower than 15 °C.

1. Fill up the sample to 50 μl with TE buffer. Add 3 μl of 10 mg/ ml RNase A, mix and incubate for 1 h at 37 °C or O/N

at 4 °C.

2. Add 0.3 vol of 5 M NaCl to the sample.

3. Fill up the volume to 600 μl with 2% CTAB–1.4 M NaCl solution.

4. Thoroughly mix the sample by inversion. If necessary, incubate the DNA at 37 °C to fully solubilize it. Some

contaminants might stay as slimy clumps and will not dissolve.

5. Add 1 vol of chloroform and thoroughly mix the sample by careful inversion. The sample should turn milky.

6. Spin 12,000–16,000 × g for 15 min at 25 °C to separate the phases.

7. Take off the upper clear phase without disturbing the white interphase using a wide bore (HMW DNA) or normal

pipette tip and transfer to a new 1.5 ml tube.

8. Add 1 vol of isopropanol to the collected aqueous phase and thoroughly mix the sample by careful inversion.

9. Spin at 2,000–4,000 × g for 30–45 min at 25 °C for HMW DNA.

10. Carefully decant or pipet off the supernatant without disturbing the pellet.

11. Wash the pellet by adding 1 ml of 70% ethanol to the tube and invert for 15 minutes. Make sure to dislodge the

pellet to remove the salt from it.

12. Spin the tube for 5 min.

10. Wash the pellet with 1 ml of 70% ethanol by resuspending it and spin again for 5 min. Remove all liquid.

11. Wash the pellet again with 1 ml of 70% ethanol by resuspending it and spin again for 5 min.

12. Remove all liquid and briefly air-dry the pellet.

13. Resuspend the pellet in 30 μl TE buffer.

**Figure S1.** Merqury K-mer spectra plots. Merqury copy number spectrum plots for *Littorina saxatilis* genome v.2 after CANU assembly (left plot), after one round of haplotigs removal with purge_dups (middle plot) and after two rounds of haplotigs removal with purge_dups (right plot). Each plot tracks the multiplicity of each k-mer found in the Illumina read set and colors it by the number of times it is found in a given assembly.

For example, k-mers found only one time in the assembly are represented by the red line that forms 2 peaks of multiplicity. The first peak represents 1-copy (heterozygous) k-mers in the genome, and the second peak represents 2-copy k-mers originating from homozygous sequence or haplotype-specific duplications. Depth of sequencing coverage determines where these peaks appear. The assembly k-mers absent from the read set are plotted as a bar at zero multiplicity, colored by the copy numbers found in the assembly. Missing single copy k-mers (read-only) are plotted in black, and k-mers present more than one time in the assembly are represented in blue, green, purple and orange.

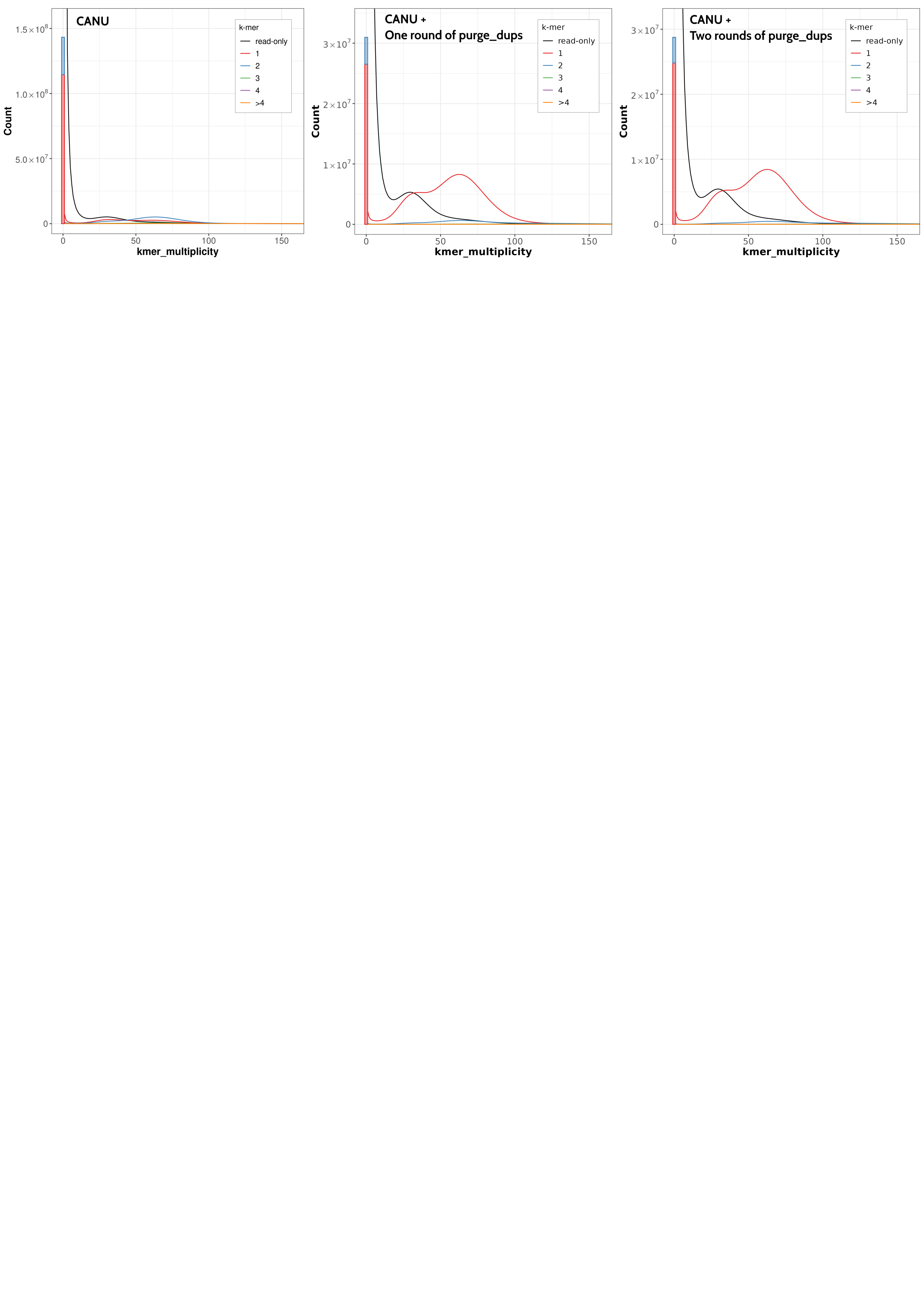

**Figure S2.** Hi-C contact map of the new Littorina saxatilis assembly, visualised in HiGlass. Chromosomes are arranged in size order from left to right and top to bottom. The left Hi-C contact map is the map obtained after YaHS scaffolding. The right map is the map obtained after YaHS scaffolding and manual curation.

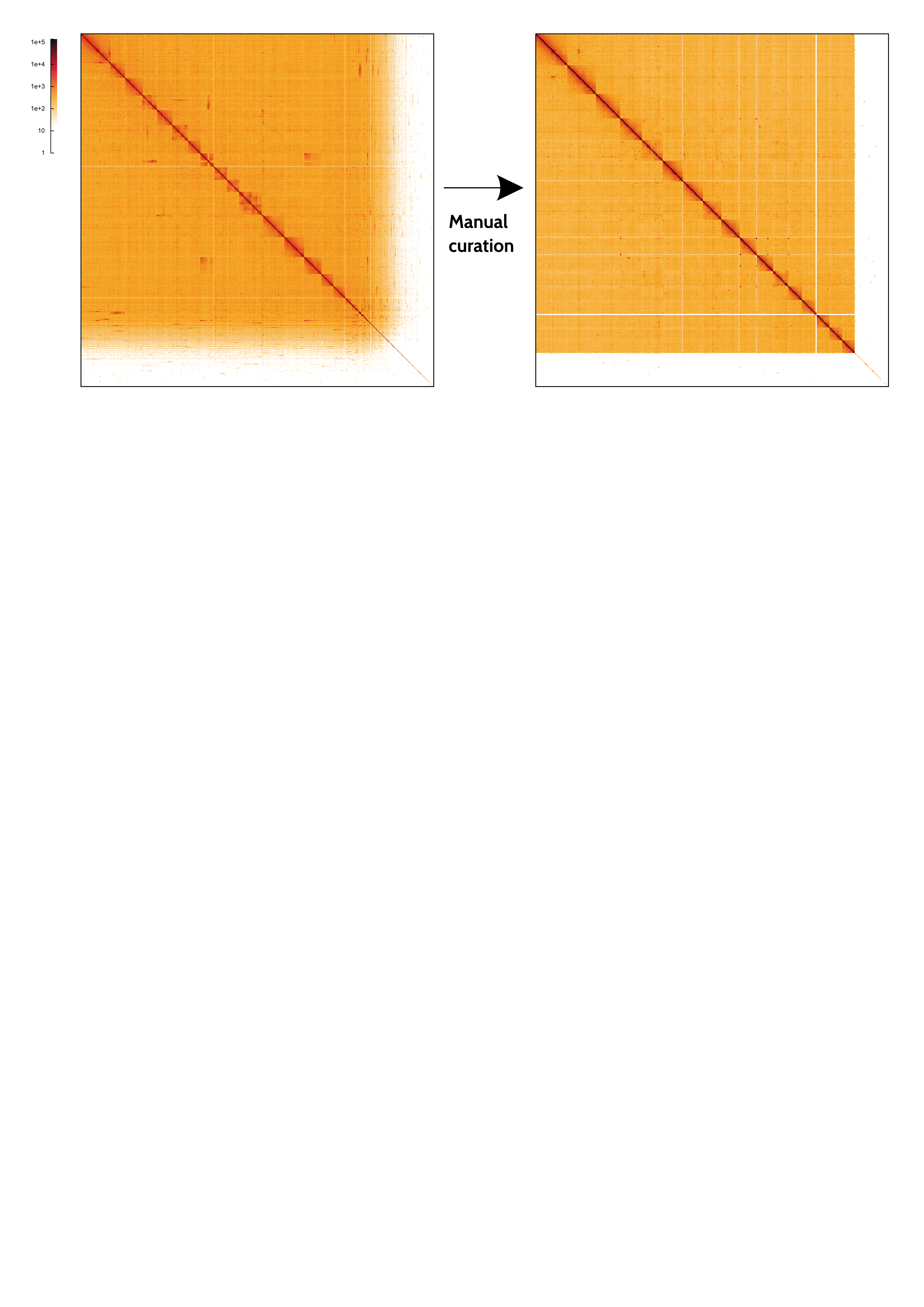

**Figure S3.** BlobToolKit GC-coverage plot. Scaffolds are coloured by phylum. Circles are sized in proportion to scaffold length. Histograms show the distribution of scaffold length sum along each axis.

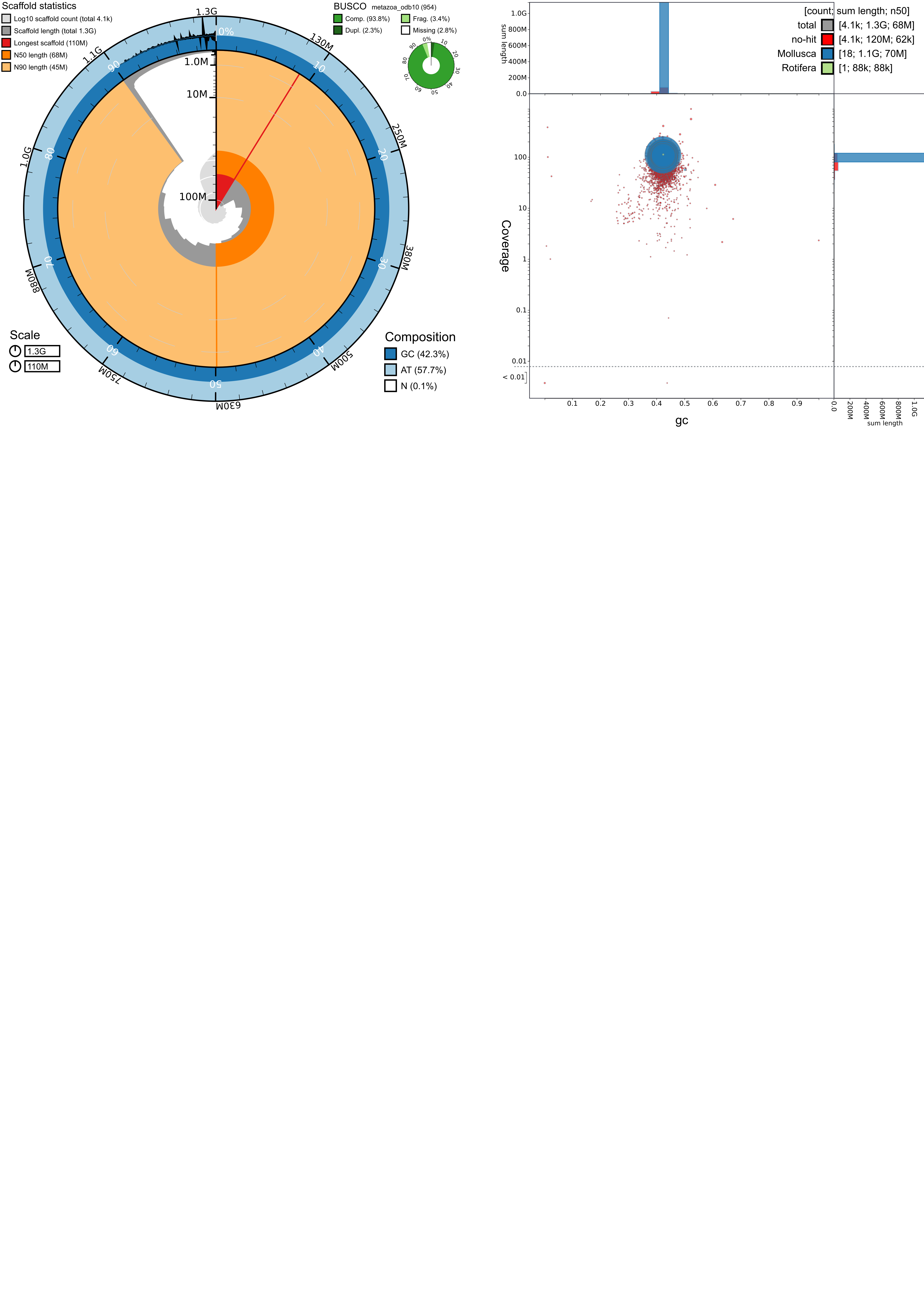

**Table S2.** Correspondence between *L. saxatilis* genome v.1 linkage groups and *L. saxatilis* genome v.2 super-scaffolds.

| ***L. saxatilis* genome v.1 linkage group** | ***L. saxatilis* genome v.2 super-scaffold** |
| --- | --- |
| **LG1** | **SUPER_1** |
| **LG2** | **SUPER_2** |
| **LG3** | **SUPER_4** |
| **LG4** | **SUPER_7** |
| **LG5** | **SUPER_9** |
| **LG6** | **SUPER_11** |
| **LG7** | **SUPER_10** |
| **LG8** | **SUPER_5** |
| **LG9** | **SUPER_8** |
| **LG10** | **SUPER_14** |
| **LG11** | **SUPER_16** |
| **LG12** | **SUPER_3** |
| **LG13** | **SUPER_17** |
| **LG14** | **SUPER_15** |
| **LG15** | **SUPER_13** |
| **LG16** | **SUPER_12** |
| **LG17** | **SUPER_6** |
